# Restoration of 3-D structure of insect flight muscle from a rotationally averaged 2-D X-ray diffraction pattern

**DOI:** 10.1101/2024.09.04.611338

**Authors:** Hiroyuki Iwamoto

## Abstract

The contractile machinery of muscle, especially that of skeletal muscle, has a very regular array of contractile protein filaments, and gives rise to a complex and informative diffraction pattern when irradiated with X-rays. However, analyzing these diffraction patterns is often challenging because: (1) only rotationally averaged diffraction patterns can be obtained, resulting in a substantial loss of information, and (2) the contractile machinery contains two different sets of protein filaments (actin and myosin) with different helical symmetries. The reflections originating from them often overlap. These problems may be solved if the real-space 3-D structure of the contractile machinery is directly calculated from the diffraction pattern. Here we demonstrate that, by using the conventional phase-retrieval algorithm (hybrid input-output), the real-space 3-D structure of the contractile machinery can be effectively restored from a single rotationally averaged 2-D diffraction pattern. In this calculation, we used an in-silico model of insect flight muscle, which is known for its highly regular structure. We also extended this technique to an experimentally recorded muscle diffraction pattern.

## Introduction

Because of their short wavelengths, X-rays are potentially capable of observing various types of samples (crystalline and non-crystalline) at an atomic resolution. Because of their high penetrability, the observation technique using X-rays is applicable to thick specimens not transparent to visible light or electron beams. This technique is even applicable to live cells or tissues of living organisms.

The drawback is that it is very difficult to fabricate X-ray lenses that ensure a high resolution. For this reason, it is common practice to directly analyze the X-rays scattered or diffracted by the samples. A problem here is that both the amplitude and the phase information of the scattered beam are required to restore the structural information of the samples, but the phase information is lost when the scattering is recorded on a detector. This is called the phase problem, and has been the largest obstacle in applying the X-ray technique to a wide range of specimens.

Recently, however, it was theoretically shown (Bates, 1982; Miao et al., 1998) and experimentally demonstrated (Miao et al., 1999) that the once-lost phase information can be recovered, meaning that a magnified image of the sample is obtained through iterative calculations, if certain conditions are met during data recording. The conditions include (1) that the X-ray beams are coherent enough to cover the entire sample, and (2) that the data sampling is finer than the Nyquist spacing, which is the inverse of the sampling size of the specimen (Miao and Sayre, 2000). The latter is commonly called the oversampling condition. Since the first experimental demonstration (Miao et al., 1999), this phase-retrieval technique and its variations have been widely employed. Today, they are called coherent diffractive imaging (CDI). CDI works very well for isolated particles with high contrast in electron density, such as metal nanoparticles. The original CDI is 2-dimensional (2-D), meaning that a 2-D image of the sample is restored from a 2-D X-ray scattering pattern.

The aim of the present work is to test whether CDI can restore the 3-D structure of a complex biological system (specifically, the sarcomeric structure of muscle) from its 2-D X-ray diffraction pattern. Such a biological system is a challenge to CDI in many ways: (1) It is an assembly of proteins in water, and has much lower density contrast compared to a metal nanoparticle. (2) It consists of fibers with helical symmetries. CDI is not very suitable for such a structure. (3) It can generate only rotationally averaged diffraction patterns, from which substantial information has been lost. (4) Expanding CDI to 3-D requires substantial computational resources and time.

The present paper demonstrates that, after appropriate expansions of the original 2-D CDI, in principle, the 3-D structure of sarcomere can be restored reasonable well from its rotationally averaged 2-D diffraction pattern. The details of the expansions are described in the following sections.

### Procedures of 3-D reconstruction

#### Structure of a sarcomere in muscle

Figure 1A is a schematic diagram of a sarcomere in insect flight muscle. A sarcomere is the minimal functional unit of muscle, and is approximately 2 x 2 µm in length and diameter. A sarcomere contains two sets of protein filaments—actin and myosin— which slide past each other to cause muscle contraction. Both filaments have their own helical symmetry. Figure 1B shows a diffraction pattern recorded from the flight muscle of a giant waterbug, *Lethocerus deyrolei*. Only a rotationally averaged 2-D diffraction pattern can be recorded, because the myofibrils that constitute a flight muscle cell are randomly oriented in rotation. Our goal is to restore a 3-D structure as shown in Fig. 1A from a 2-D diffraction pattern as shown in Fig. 1B. Insect flight muscles are known to have a very regular structure, and the symmetries of the filaments are well documented (Tregear et al., 1998, 2004). Therefore, they are well-suited for modeling in this type of study.

**Figure 1.**
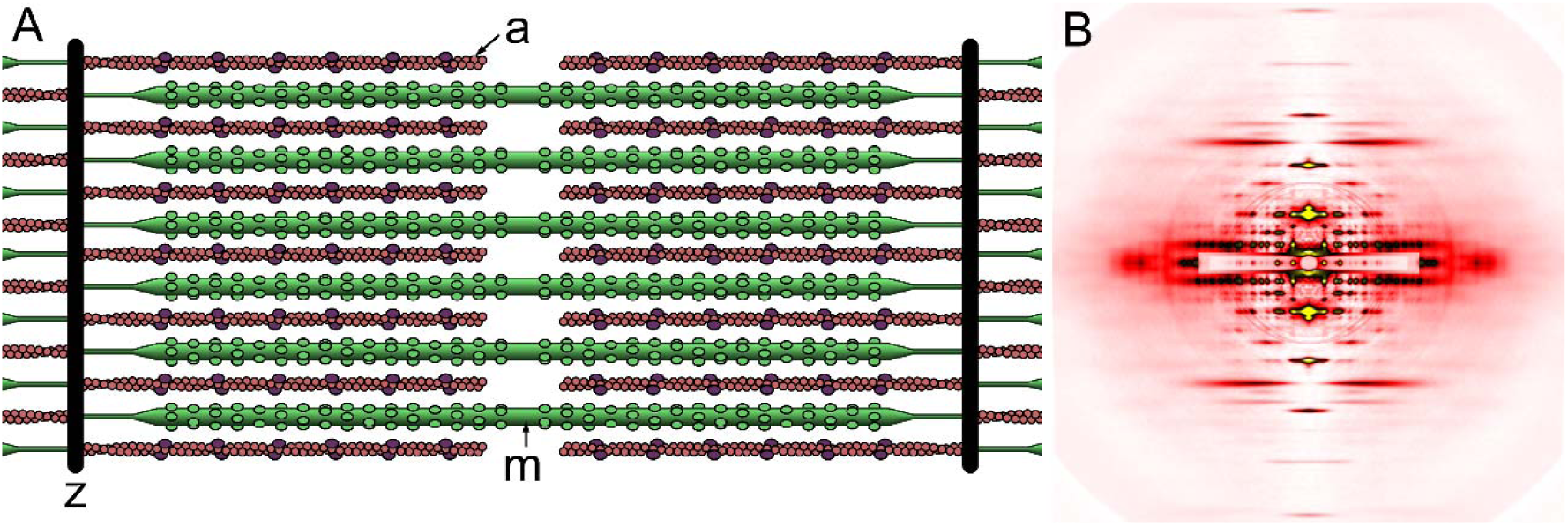
Structure of a sarcomere in insect flight muscle and an X-ray diffraction pattern recorded from it. (A) schematic diagram of sarcomeric structure; (B) diffraction pattern recorded from the flight muscle of a giant waterbug, *Lethocerus deyrolei*. a, actin filament; m, myosin filament; z, Z-membrane that separates two neighboring sarcomeres. The diagram in (A) is adapted from Iwamoto, 2011.

#### Principle of 2-D CDI

The principle of CDI in the original form has been documented in many references (e.g., Miao and Sayre, 2000), and it is only briefly reiterated here.

A diffraction pattern recorded from a sample has amplitudes proportional to the square of the structure factor (Fig. 2A), which is the Fourier transform of the sample’s electron densities and a function in reciprocal space. Initially, the structure factor lacks phase information, so a random phase is assigned. Applying inverse Fourier transformation to the structure factor reconstructs the sample’s image in real space. Due to the incorrect phase, an inaccurate image of the sample is generated (Fig. 2B), showing densities outside the maximal area in which the sample is expected to exist. Then the densities outside this area (called support) are deleted (Fig. 2C). This process is called real-space correction. After this, the image is subjected to Fourier transform to obtain the structure factor again (Fig. 2D). The structure factor now contains phase information that is expected to be closer to the correct value than the initial random phase. After its amplitude is scaled to the observed value (reciprocal-space correction, Fig. 2E), the entire process is repeated. During the iterations, the error in the phase is gradually reduced, and finally the correct image of the sample is restored. Recent calculation procedures incorporate a process to automatically reduce the support to fit the contour of the image, resulting in better convergence (the shrink-wrap algorithm, Marchesini et al., 2003).

**Figure 2.**
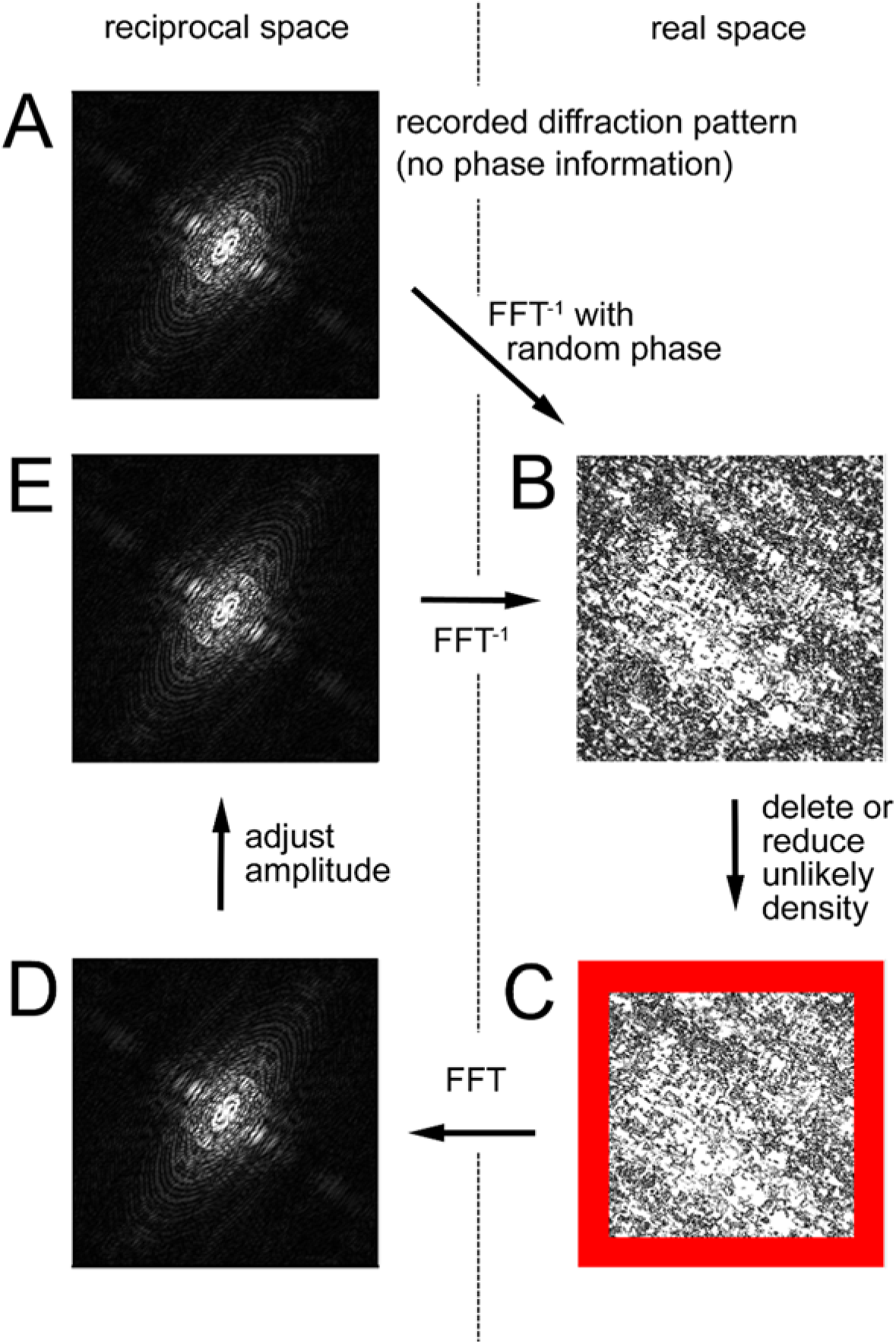
Principle of 2-D CDI. The recorded structure factor (A) is given a random phase. It is then subjected to inverse Fourier transformation to generate a real-space image (B). After unlikely densities are corrected (C), the image is subjected to Fourier transformation to obtain a structure factor (D). After its amplitude is rescaled to the observed values (E), it is subjected to inverse Fourier transformation again (B). The calculations are iterated in the sequence B-C-D-E until the error is minimized (convergence). In (C), the red area is outside the support.

The example shown in Fig. 2 is taken from an in-silico experiment using a logo mark of SPring-8. For such an isolated, high-contrast object, CDI performs well and calculations usually converge within 400-600 iterations (Fig. 3).

**Figure 3.**
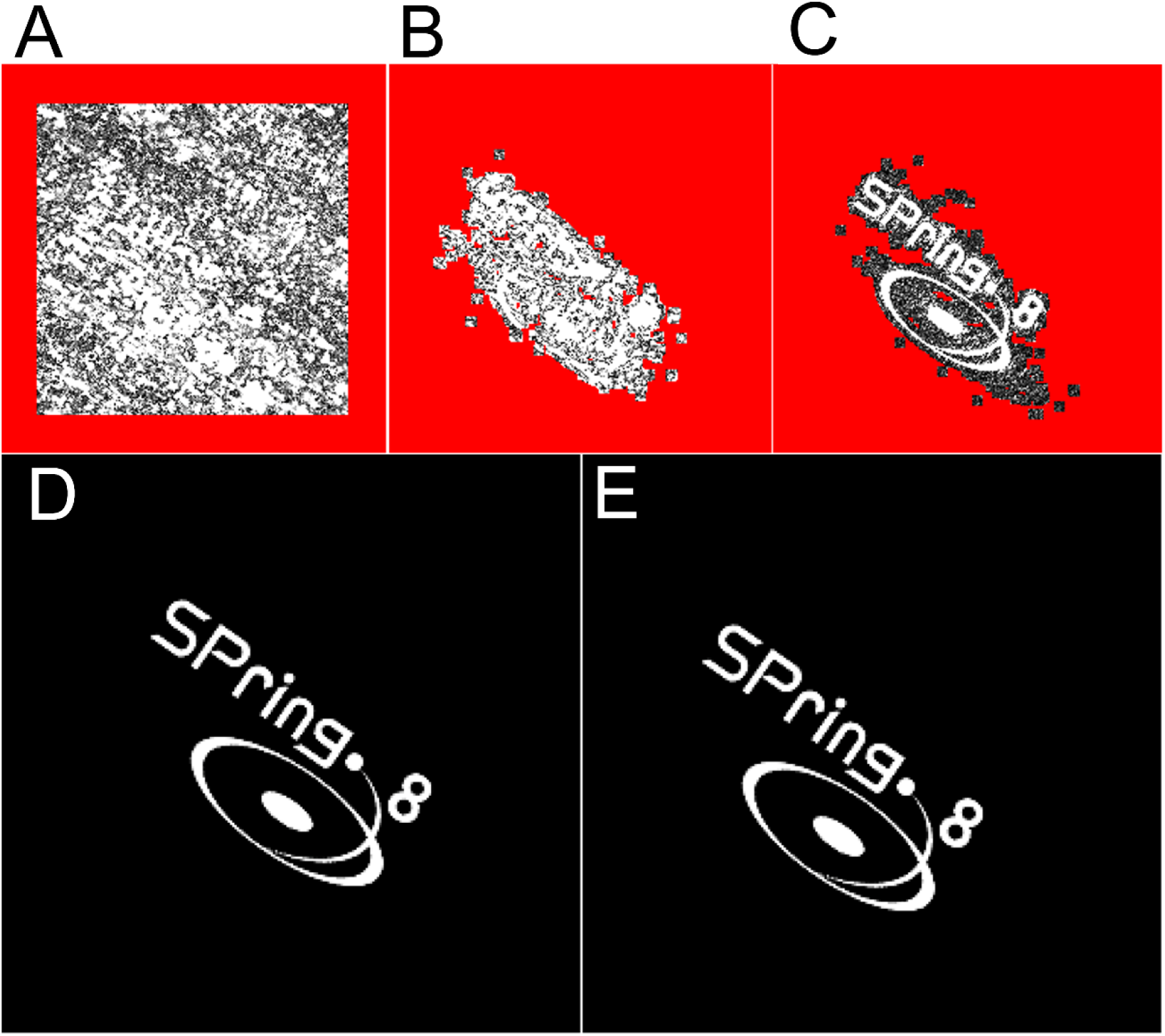
In silico demonstration of 2-D CDI by using the SPring-8 logo as a sample. (A) support area immediately after the start of iterations (the red area is outside the support). (B) support area reduced by the shrink-wrap algorithm at 250 iterations. (C) status of calculation at 290 iterations when the calculation started to converge. (D) correct answer; (E) reproduced image after 600 iterations.

#### 2-D CDI does not perform well with fibrous objects

On the other hand, CDI is unsuitable for fibrous materials with periodic structures, which are commonly found in living organisms. Examples shown in Fig. 4 are a row of the letter ‘a’ (Fig. 4A) and the 2-D projection of an actin filament, decorated with the head parts enzymatically isolated from myosin molecules (decorated actin, Fig. 4B). The entire structure follows the helical symmetry of the actin filament (28 monomers in 13 turns). When CDI is applied to the image of decorated actin, the calculation does not converge even after thousands of iterations, and the restored image is not satisfactory (the right images in Fig. 4A and B). One reason why CDI does not work for fibrous samples may be that the scattering pattern is separated into discrete layer lines (Fig. 4C, D), preventing the oversampling condition from being met.

**Figure 4.**
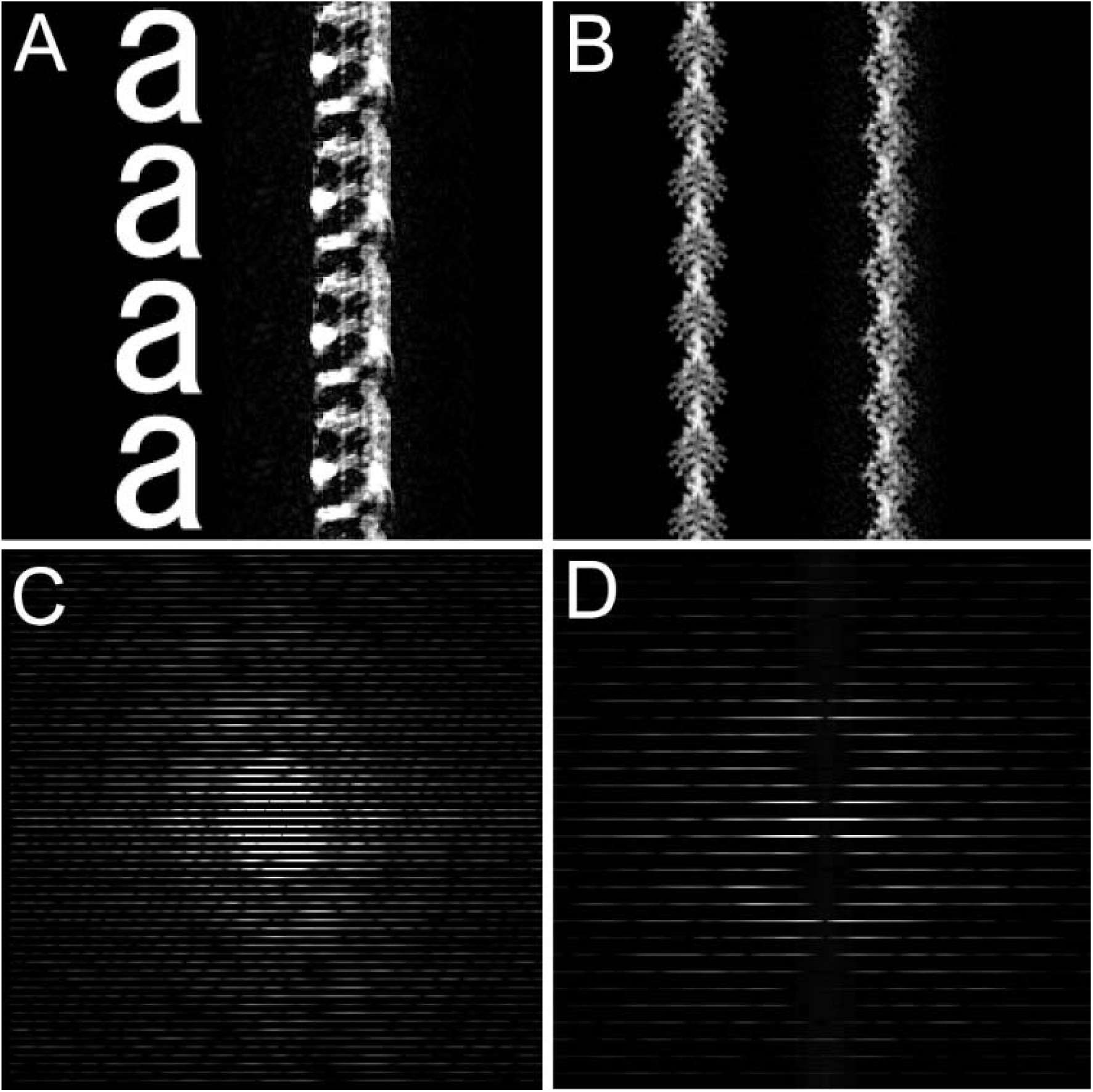
2-D CDI applied to fibrous materials with periodic structures (in silico experiments). (A) a vertical row of the letter ‘a’. The right image is the unsuccessful result of CDI calculation (the calculation never converges). (B) 2-D projection of decorated actin (see text for details). The result of the CDI calculation is, again, unsuccessful. (C) and (D) are the structure factors calculated from the left-hand images in (A) and (B), respectively. The 3-D coordinate data of decorated actin was taken from Lorenz et al. (1993).

#### Expansion to 3-D CDI

Then we consider expanding the 2-D CDI to the 3-D CDI. Mathematically, one can achieve this by simply replacing 2-D Fourier transformation with 3-D Fourier transformation. In real experiments, one must collect a large number of diffraction patterns while rotating the sample and combine them into a single 3-D diffraction pattern, in a procedure similar to X-ray computed tomography. For real biological samples, it is difficult to collect a large number of diffraction patterns due to radiation damage, but conducting in-silico experiments to evaluate 3-D CDI is much easier. If the object is an isolated particle, the calculation converges without any problems. The question is whether a fibrous object can be restored using 3-D CDI.

The examples used to test 3-D CDI are the 3-D models of decorated actin (Fig. 5A) and an axoneme of eukaryotic flagella (Fig. 5B). When these objects are subjected to the same procedure shown in Fig. 2, except that the calculation is carried out in a 3-D space, the calculation converges in both cases, and the model structures are perfectly restored. As shown in Fig. 5, restoring a short segment of the filament using a short support seems to result in better convergence. The reason why a fibrous object can be restored using 3-D CDI is unclear, but it may be due to the oversampling condition being met along two axes (X and Y), whereas in 2-D CDI, it is met along only one axis (X).

**Figure 5.**
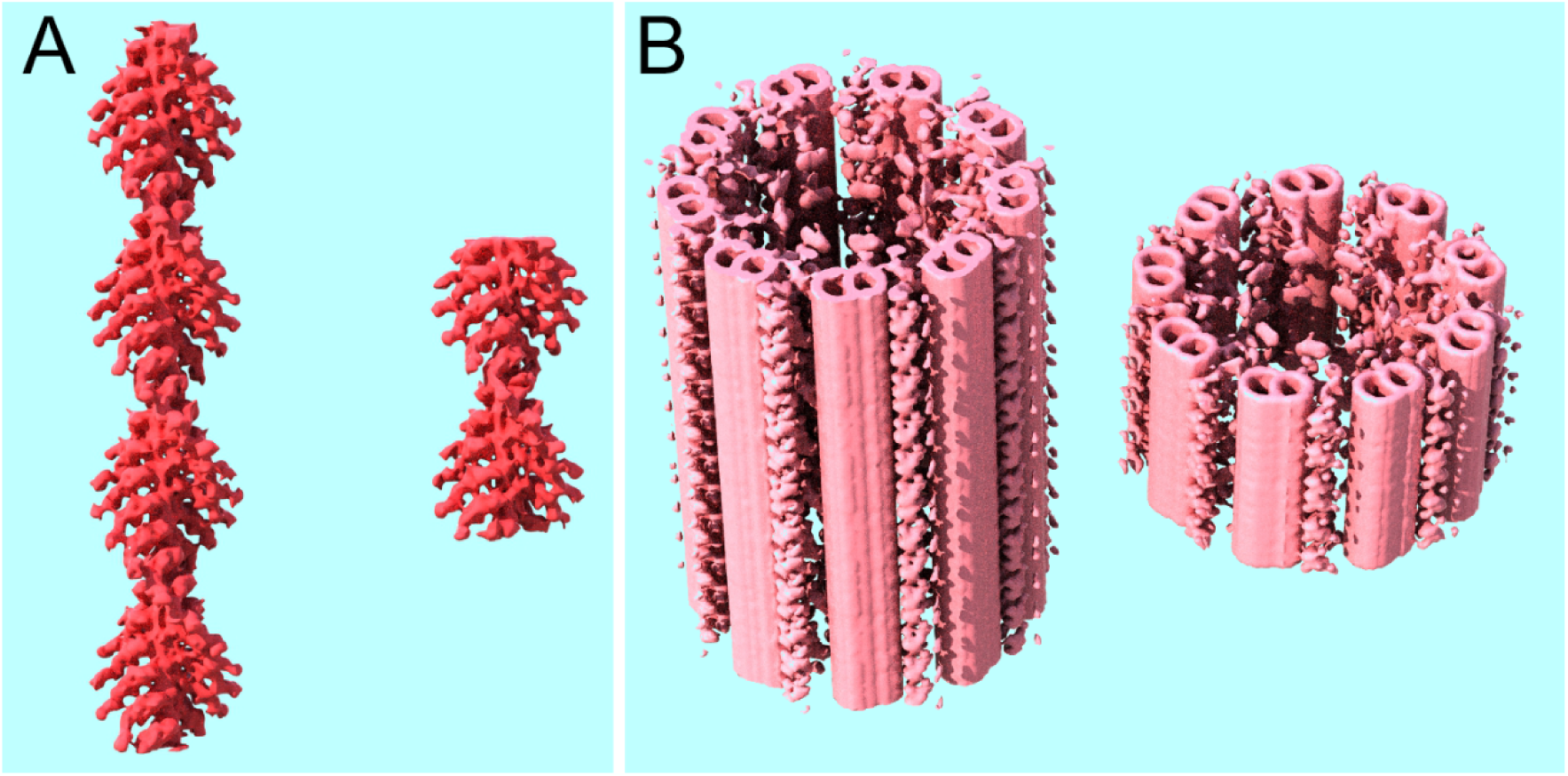
Demonstration that 3-D CDI works for fibrous materials with periodic structures. (A) 3-D model of decorated actin; (B) 3-D model of an axoneme of eukaryotic flagella. In each, the left image is the surface-rendered view of the model and the right image is that of the restored structure. The coordinate data of decorated actin are identical to those in Fig. 4. The density distribution of the axoneme was constructed from the tomogram provided by Bui et al. (2012).

#### Restoration of the 3-D sarcomeric structure

Having established that 3-D CDI works for fibrous objects, we proceed to the final objective: restoring the 3-D structure of the sarcomere of insect flight muscle from its rotationally averaged 2-D diffraction pattern. Details of the procedure of calculations are described below.

First, the basic lattice structure of actin and myosin filaments and the helical symmetry of both filaments are well documented (e.g., Tregear et al., 1998, 2004), and we make full use of this information. As for the lattice structure, the filaments form a hexagonal lattice, and a single actin filament is located midway between two neighboring myosin filaments (Fig. 6A). The unit cell size (*d*1,0 spacing) is approximately 45 nm. As for axial periodicities, myosin filament has a structure of 4-start helix, which has 16 monomers in 2.5 turns with a monomer interval of 14.5 nm. Therefore, the basic periodicity of the myosin filament is 232 nm (14.5 x 16 = 232 nm; Tregear et al., 1998, 2004; Fig. 6B). The actin filament has a structure of a 1-start helix, with 28 monomers in 13 turns and a monomer interval of 2.75 nm. Therefore, the basic periodicity of the actin filament is 77 nm (2.75 x 28 = 77 nm; Tregear et al., 1998, 2004), but if it is multiplied by 3, it is also approximately 232 nm (Fig. 6B). Thus, the entire system can be regarded as having a basic axial repeat of 232 nm.

**Figure 6.**
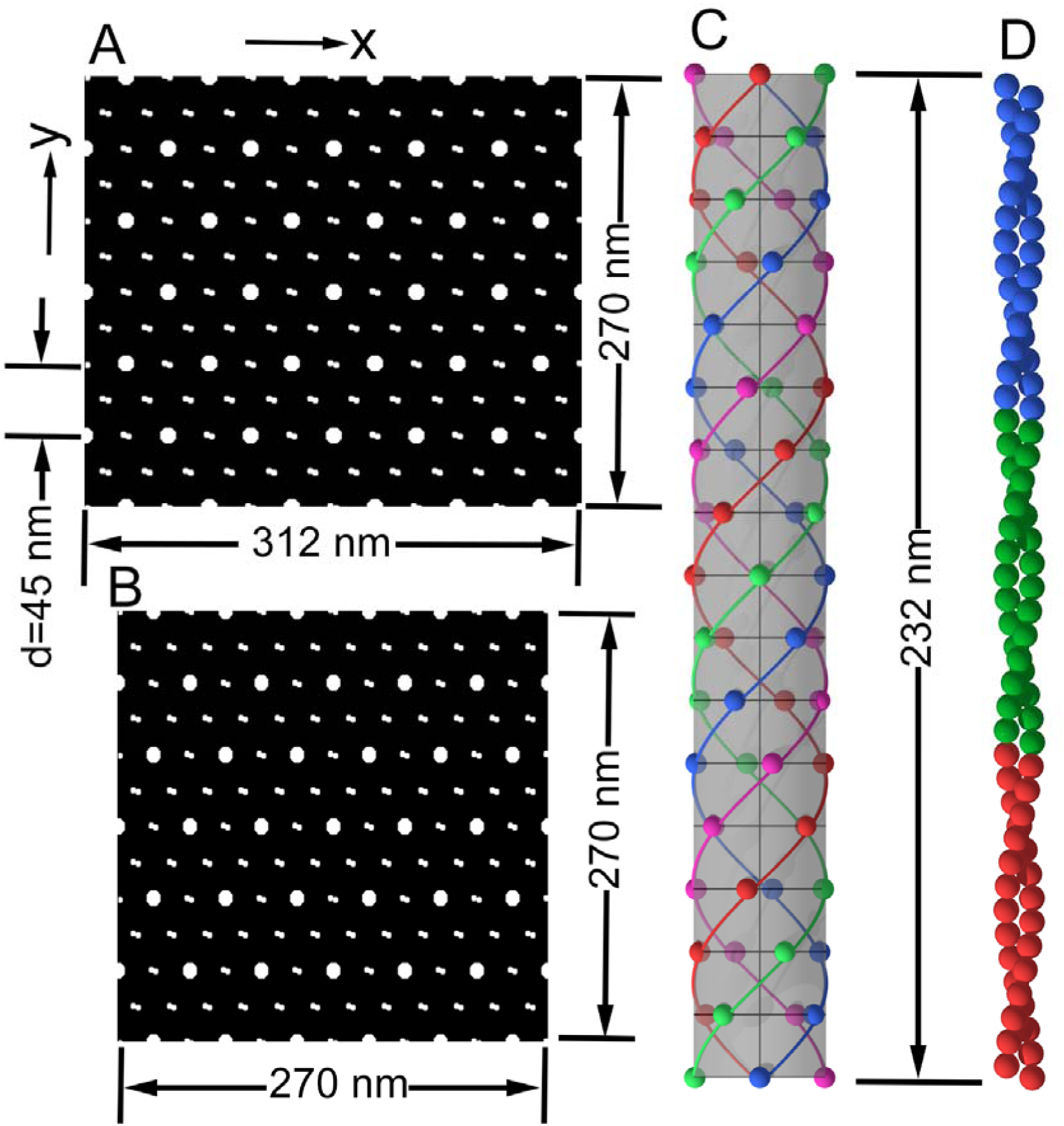
Optimization of the model of sarcomeric structure for fast Fourier transformation (FFT). In order for FFT to work properly, the 3-D calculation space should be the size 2^n^ x 2^n^ x 2^n^ (n is an integer) and it should contain integer numbers of repeating units in all of X, Y, and Z directions. When the d-spacing of the unit cell is 45 nm, an area of 270 x 312 nm in the X-Y plane (A, across the axes of myosin and actin filaments) can contain integer numbers of repeating units in both X and Y directions. This area is compressed to 270 x 270 nm (B). In the Z direction, both myosin (C) and actin (D) filaments have a periodicity of 232 nm, and they are extended to 270 nm. In this way, the sarcomeric model can be accommodated in a 2^n^ x 2^n^ x 2^n^ cube. In (A) and (B), the larger circles represent myosin filaments, and the smaller circles represent actin filaments.

To ensure reasonably short calculation time, the iterative Fourier and inverse Fourier transformations must be performed using the fast Fourier transform (FFT) algorithm. To implement this, the data space to be processed must be a cube, with its side length of 2^n^ voxels (e.g. 128, 256, etc). In the following calculation, the size of 256 was used. To avoid boundary problems, the cube must contain an integral number of repeating units. Here, in the Y direction (X and Y axes are perpendicular to the axis of the filaments), 6 unit cells of the hexagonal lattice are accommodated (side length = 45 x 6 = 270 nm). In the X direction, it is impossible to accommodate an integral number of repeating units. Therefore, the filament lattice was compressed to √3/2 so that 6 myosin-myosin intervals are accommodated in the side length of 270 nm. In the Z direction (along the filament axis), the 232-nm repeat was expanded to 270 nm. In this system, the lowest-order prominent layer-line reflection at *d* = 38.7 nm (the 6th order of the 232-nm repeat) appears at 6 voxels from the equator.

The model structure of the sarcomere constructed in this way is shown in Fig. 7A. From this structure, a rotationally averaged 2-D structure factor can be calculated (Fig. 7B). This 2-D structure factor is used as the observed data in the following 3-D CDI calculation.

**Figure 7.**
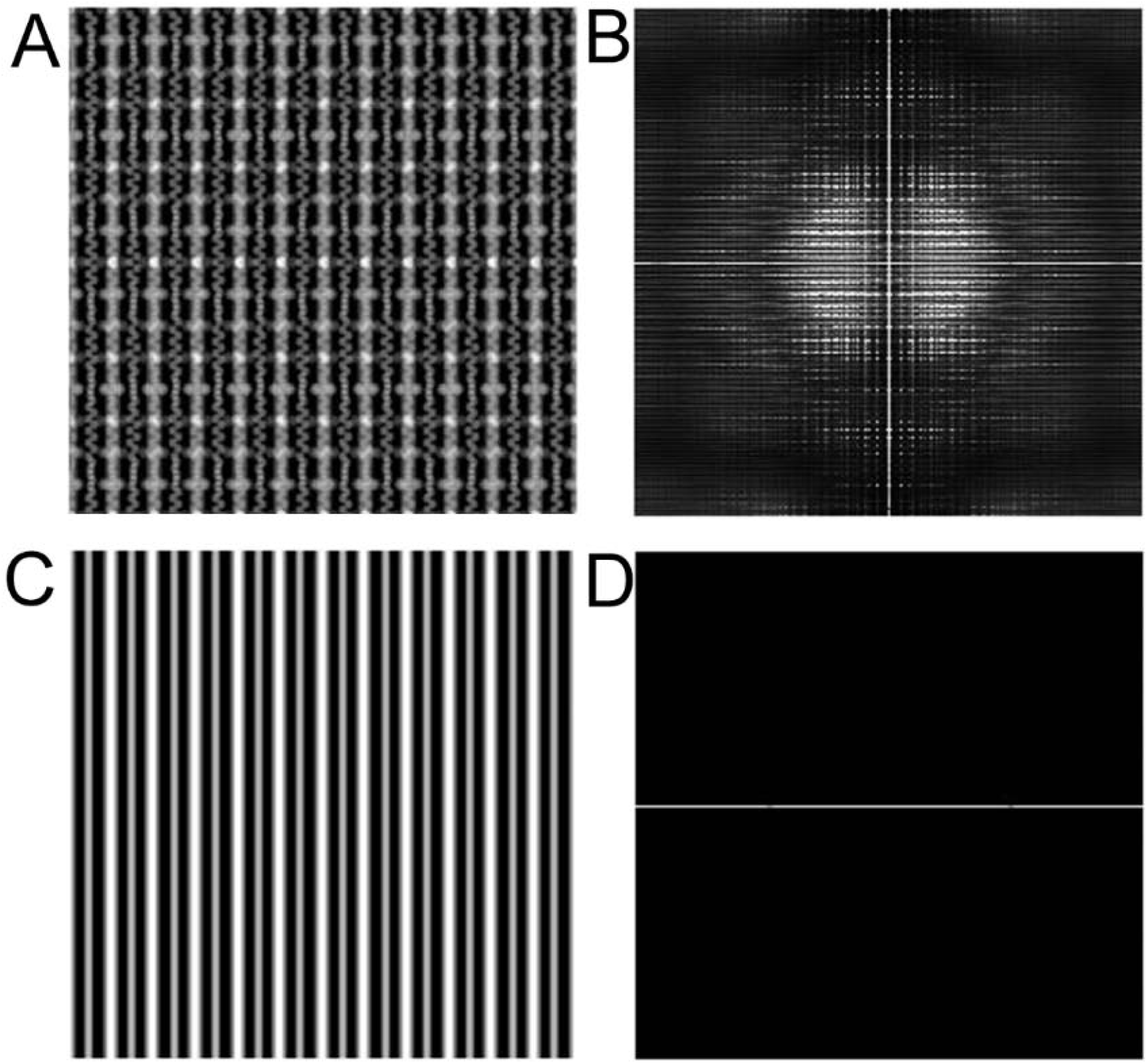
Models of sarcomeric structures of insect flight muscle. (A) side view of the 3-D model built according to the procedure shown in Fig. 6, using the known arrangement of the hexagonal lattice of actin and myosin filaments, and the known helical symmetries of individual actin and myosin filaments. The space is 256 x 256 x 256 voxels. (B) rotationally averaged 2-D structure factor calculated from (A). This pattern is used as the observed structure factor, without phase information, in the following 3-D CDI reconstruction. (C) side view of the starting model structure. It has an arrangement of actin and myosin filaments in a hexagonal lattice identical to that in (A). The individual actin and myosin filaments are expressed as featureless cylinders. (D) rotationally averaged 2-D structure factor calculated from (C), containing only the equatorial reflections. In the following 3-D CDI calculations, the structure in (C) will be modified to make its structure factor approach that in (B).

The 3-D CDI calculation begins with a starting structure shown in Fig. 7C. In this structure, actin and myosin filaments are correctly positioned within the lattice, but they are featureless cylinders without axial periodicities. The rotationally averaged 2-D structure factor of this initial structure shows only equatorial reflections (Fig. 7D). We will test whether the periodic structure shown in Fig. 6A is restored in the initial structure after cycles of 3-D CDI calculations.

Compared with the simple 3-D CDI explained in the previous section, the restoration of the 3-D structure from a rotationally averaged 2-D structure factor involves two additional steps in reciprocal space: rotational averaging of the 3-D structure factor (Fig. 8, D to E) and the restoration of the 3-D structure factor from the rotationally averaged 2-D structure factor (Fig. 8, F to G). The former step is easy, but the latter requires careful consideration. Here, we adopted a procedure used by Grant (2018) to restore the 3-D structure of proteins from their 1-D solution scattering data. In 3-D reciprocal space, he scaled all the data at a specific radius from the origin to the observed amplitude. Likewise, we scaled all the data at a specific radius from the Z axis to the observed amplitude. This correction was made within a single X-Y plane and repeated for all Z levels.

**Figure 8.**
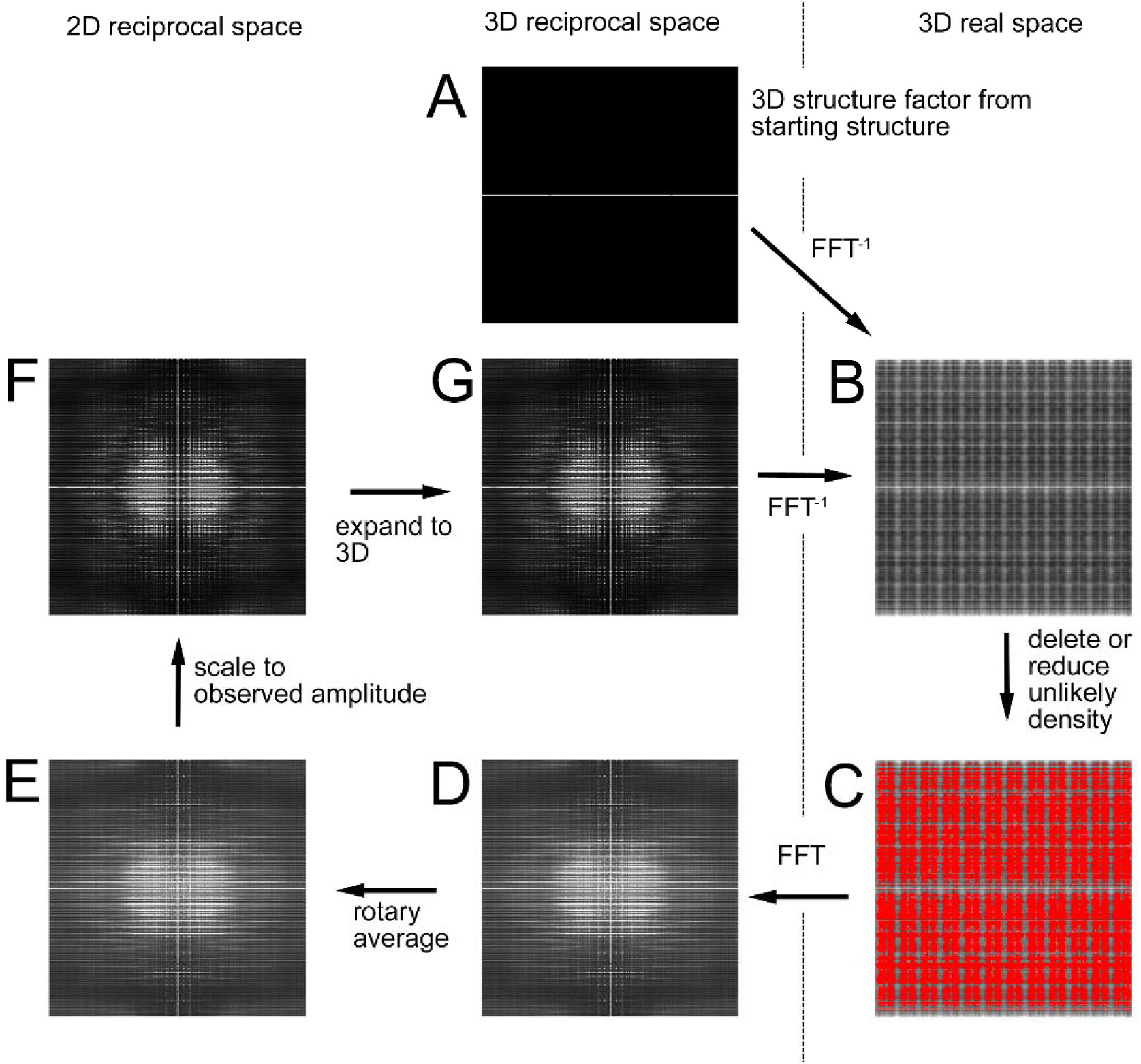
Procedure for 3-D CDI calculations involving rotationally averaged 2-D structure factor. The procedure is basically identical to that in 2-D CDI as shown in Fig. 2, except that it has two additional steps, namely, rotational averaging of the 3-D structure factor (D to E) and the expansion of the corrected 2-D structure factor to 3-D (F to G). Instead of assigning random phase information to the observed structure factor, a structure factor (A) is calculated from the starting structure (Fig. 7C), and the initial real-space data are the starting structure itself.

#### Results of 3-D restoration of model structure

Figure 9 shows the time course of the convergence of the calculation. The Fourier amplitude error (difference between observed and restored structure factors) was quickly reduced to less than 20% within approximately 10 iterations. After this, the error continued to decrease with oscillations and was reduced to 1-2% after 500 iterations. Figure 10 compares the correct and restored 2-D structure factors as well as the side views of the correct and restored lattice structures after 500 iterations. The structure factors are visually indistinguishable. The restoration of the lattice structure is not perfect, but the basic periodic structures seem to have been reasonably well restored in both actin and myosin filaments. As the error kept decreasing, an even better structure would have been restored after more than 500 iterations. The surface-rendered views of the correct, starting, and restored 3-D structures of the filament lattice are shown in Fig. 11. By using a Xeon-based workstation, the CPU time for 500 iterations was about 48 hours.

**Figure 9.**
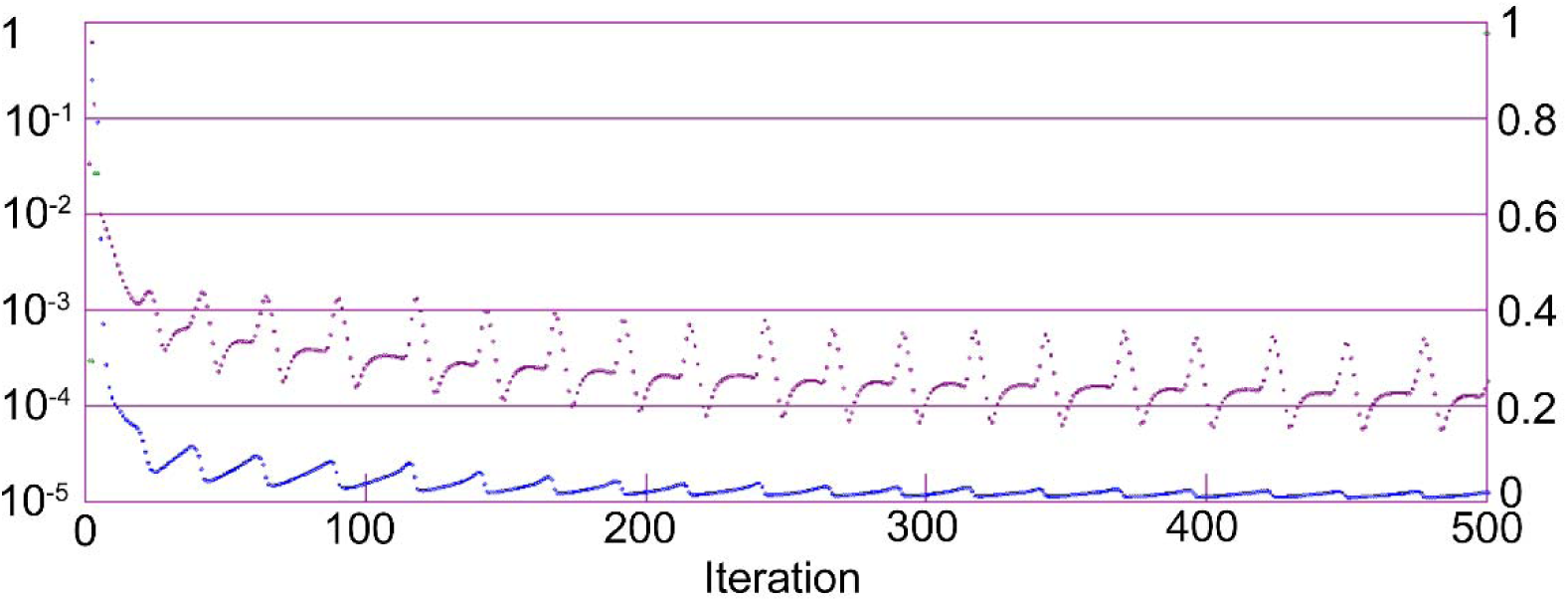
Time course of the convergence of the 3-D CDI calculation using a 2-D diffraction pattern calculated from the model structure. The Fourier amplitude error in reciprocal space (blue, linear scale on the right vertical axis) quickly decreased within ∼10 iterations, and after 500 iterations, it was 1-2%. The magenta data represent the density outside the support in real space (logarithmic scale on the left vertical axis), which was reduced to 0.0001% after 500 iterations.

**Figure 10.**
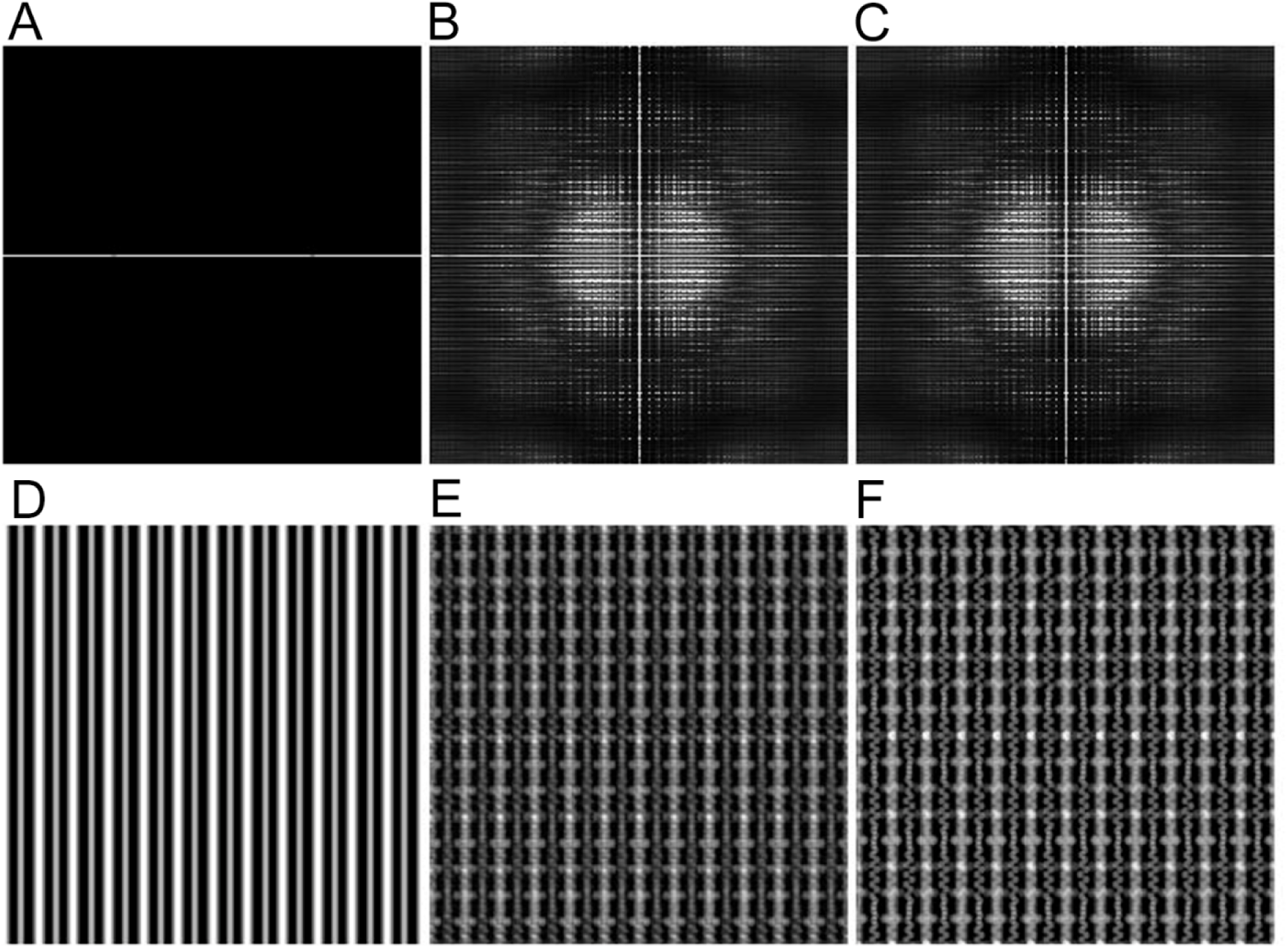
Comparison between correct and restored structure factors and the 3-D sarcomeric lattice structures. (A) rotationally averaged 2-D structure factor calculated from the starting structure (D, Fig. 7C). (B) 2-D structure factor restored after 500 iterations. (C) correct structure factor calculated from the model structure (F, Fig. 7A). (D-F) density projections of the starting structure, restored structure, and the correct model structure, respectively.

**Figure 11.**
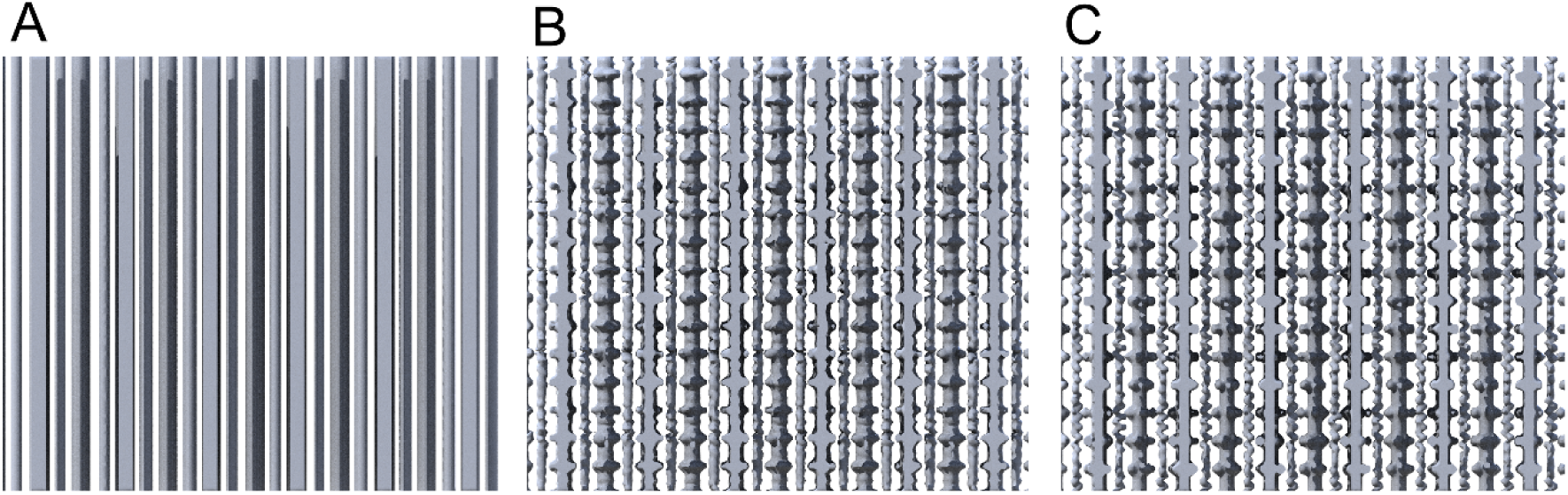
Surface-rendered views of the 3-D structures in real space. (A) starting structure; (B) structure restored after 500 iterations; (C) correct model structure.

#### Expansion to actually recorded diffraction pattern

We tested whether the procedure described above is effective for diffraction patterns actually recorded from flight muscle specimens. We used the diffraction pattern shown in Fig. 1B, recorded from *Lethocerus* flight muscle fibers. The original diffraction pattern had a size of 1000 × 1018 pixels. This was scaled down to match the size of the model diffraction pattern in Fig. 7B. To do this, we needed to expand it by 1.15 times in the equatorial direction, and then reduce the entire pattern by 0.187 times. The intensities were square-rooted to convert them into the structure factor. The periphery of the pattern had a small distortion due to the image intensifier used for recording, and it was corrected. The final size of the pattern was 256 x 256 pixels, but the area of the recorded pattern was smaller than that (Fig. 12.) The area outside the pattern was filled with a constant that approximately matched the background level of the pattern.

**Figure 12.**
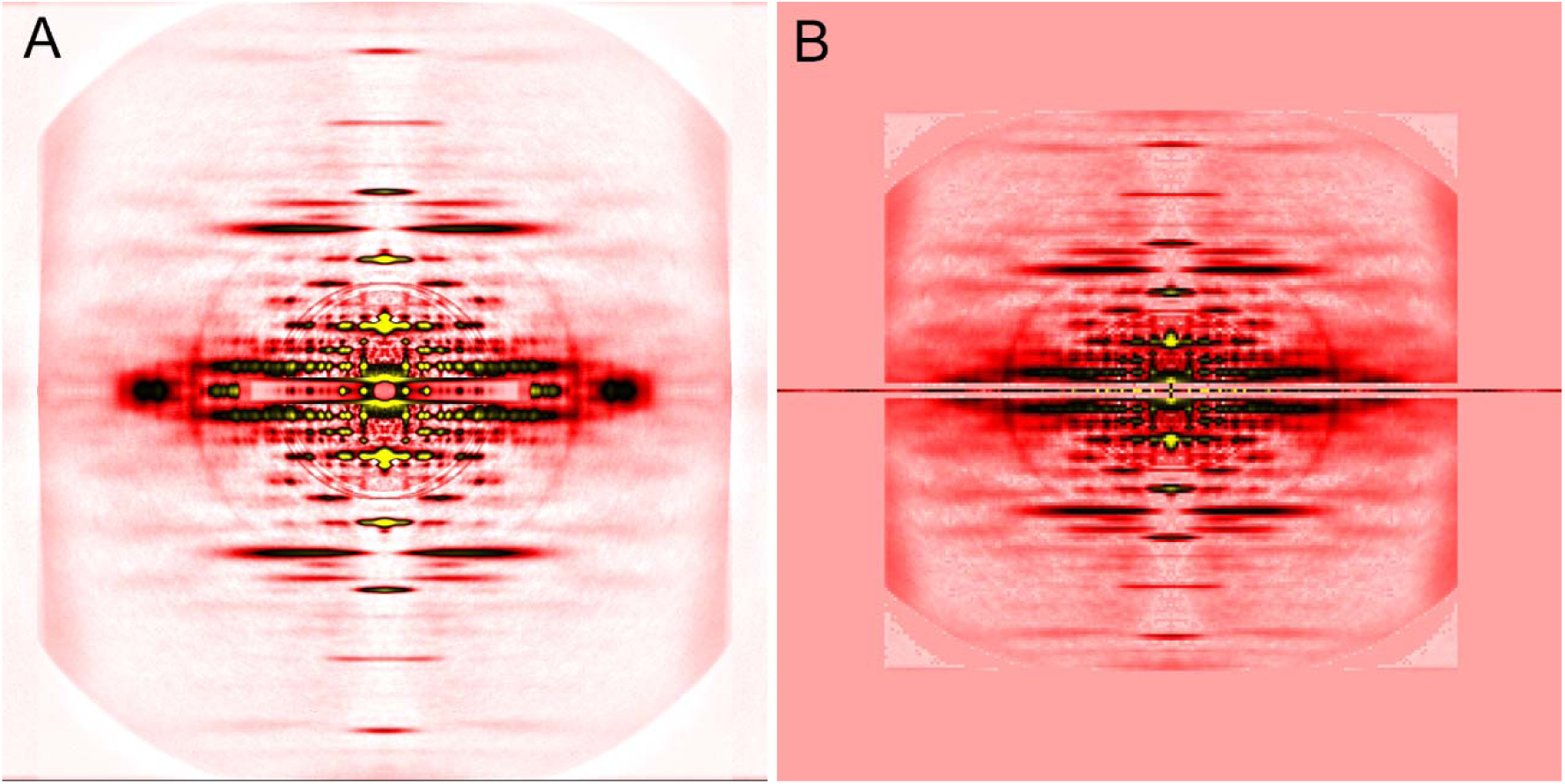
Original and processed diffraction patterns of Lethocerus flight muscle fibers. (A) full-sized diffraction pattern (1000 × 1018 pixels). The four quadrants were averaged and the background was subtracted as described by Iwamoto et al., 2013. This pattern is identical to that in Fig. 1B. (B) The same diffraction pattern processed for 3-D reconstruction. It was scaled to the same size as the pattern calculated from the model (Fig. 7B), its square root was calculated to convert to the structure factor, its equator was replaced with that of the model pattern (Fig. 7B), and the outside of the data area was filled with a constant. The whole area is 256 x 256 pixels, but its image is magnified 4x to facilitate comparisons.

When this scaled pattern was used as it is, the 3-D CDI calculation was not satisfactory. We found that precisely recorded equatorial reflections were crucial for restoring the correct myofilament lattice structure. Therefore, the equatorial reflections of the recorded pattern were replaced with those calculated from the model structure (Fig. 7B), leaving the rest of the pattern unchanged (Fig. 12B).

#### Results of 3-D restoration of real sarcomeric structure

To scale the recorded diffraction pattern, we needed to reduce its size to smaller than 256 x 256 pixels (Fig. 12B). The details in the original diffraction pattern were lost. Because of this, we did not expect that a detailed sarcomeric structure would be restored even if the calculations themselves were successful. In the original diffraction pattern, reflections up to the 1st actin meridional reflection (at 2.7 nm^-1^) were recorded, limiting the maximal spatial resolution to this value (in the model calculation, it is around 1 nm^-1^).

Figure 13 shows the time course of the convergence of the calculation. As in the calculation for the model structure, the Fourier amplitude error was quickly reduced to less than 20% within approximately 10 iterations, and after this, it was gradually reduced with occasional reversions. After 800 iterations, it was reduced to 1.8%, indicating good convergence. Figure 14 compares the restored and starting structure factors. Except for the streak on the meridian, the restored structure factor is indistinguishable from the starting structure factor.

**Figure 13.**
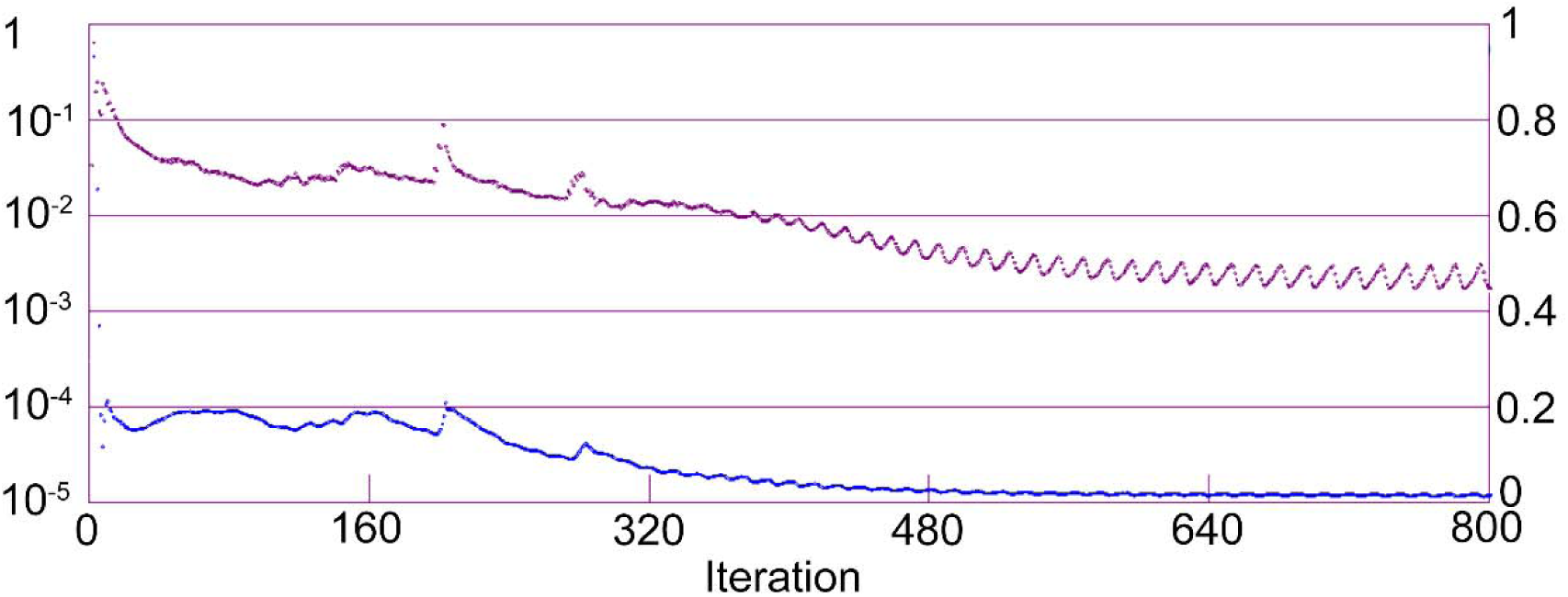
Time course of the convergence of the 3-D CDI calculation using a diffraction pattern recorded from *Lethocerus* flight muscle fibers. Plotted in the same manner as shown in Fig. 9. As in Fig. 9, the Fourier amplitude error in reciprocal space (blue) quickly decreased within ∼10 iterations, and after 800 iterations, it was 1.8%. The density outside the support in real space (magenta) was reduced to less than 0.01% after 800 iterations.

**Figure 14.**
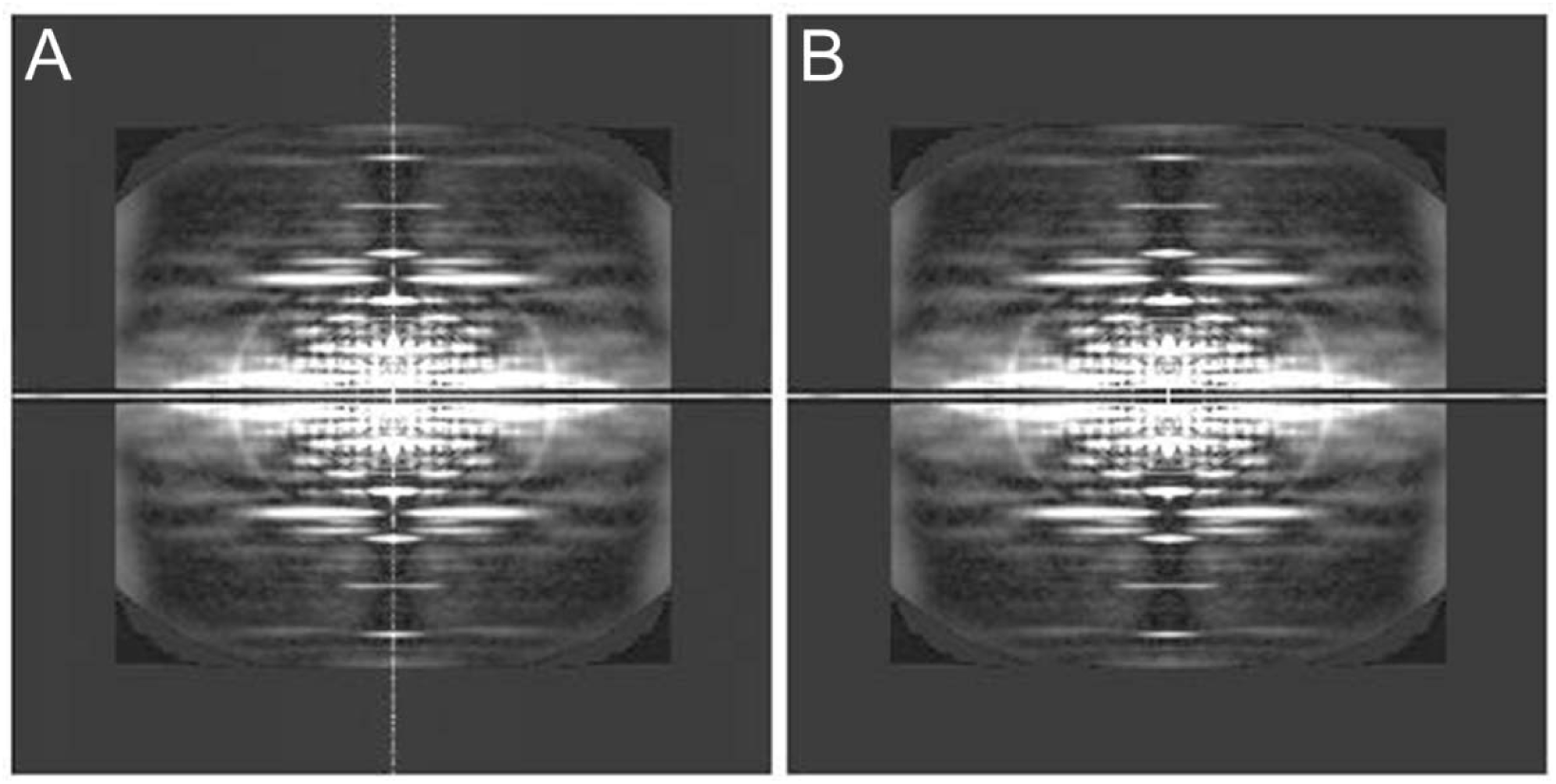
Restored (A) and starting (B) structure factors of Lethocerus flight muscle fibers. The image in (A) was obtained after 500 iterations. (B) shows the starting structure factor and is identical to Fig. 12B.

Figure 15 shows the surface-rendered view of the restored structure. The myosin filaments have bulges reminiscent of myosin heads with a 14.5-nm periodicity (blue arrowheads). Individual myosin heads are not resolved. The actin filaments often show features of individual monomers with a 5.9-nm short-pitch left-handed single-stranded helix (red arrowheads) and a 38.7-nm long-pitch double-stranded helix (magenta arrowheads).

**Figure 15.**
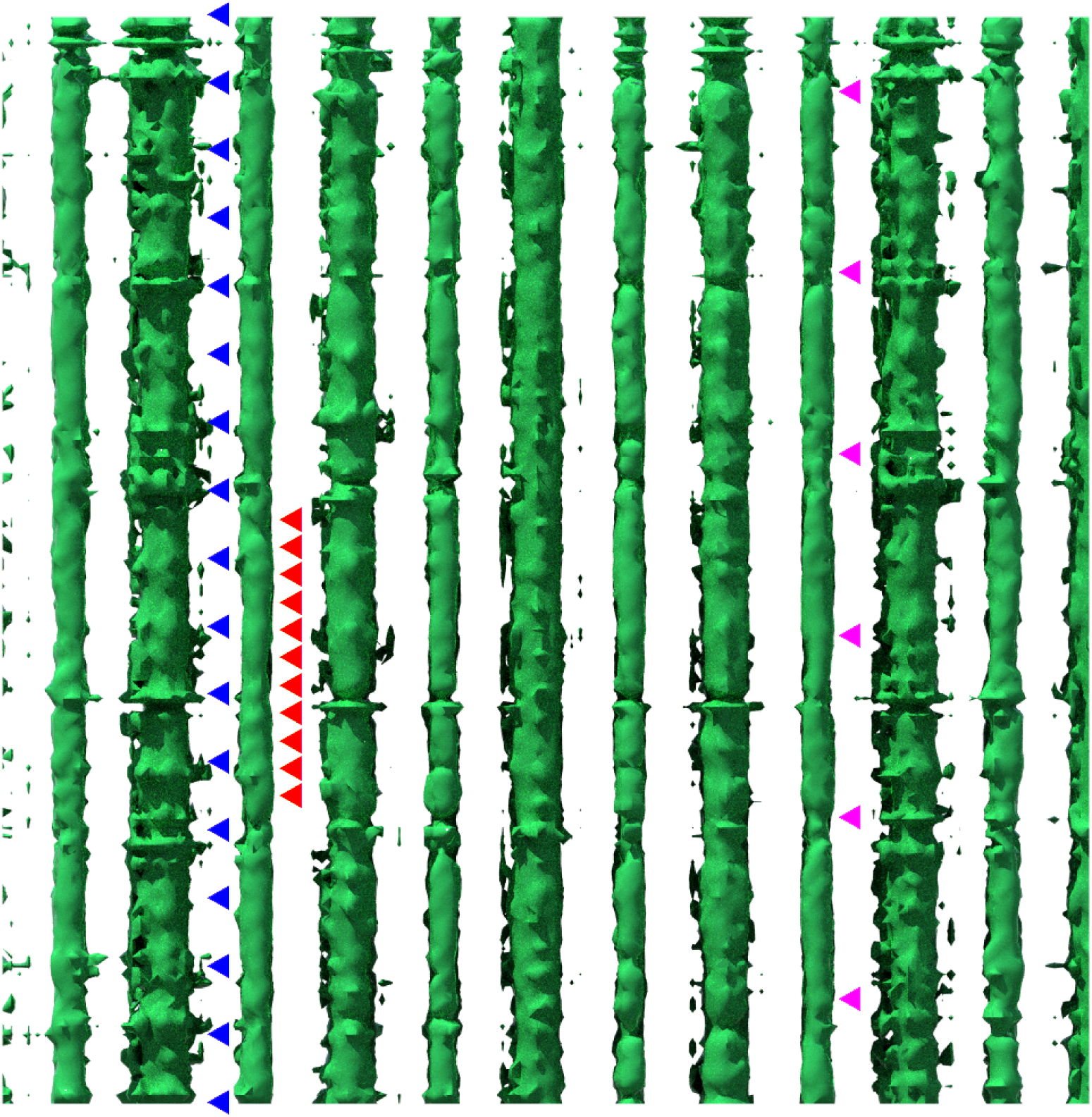
Surface-rendered view of the 3-D structure restored from the structure factor of Lethocerus flight muscle fibers. The thicker rods are myosin filaments, and the thinner ones are actin filaments. The blue, red, and magenta arrowheads indicate the spacings of myosin heads (14.5 nm), actin monomers (5.9 nm), and the cross-over repeats of the double-stranded actin helix (38.7 nm), respectively.

These periodicities are more clearly observed in the density projections of the restored structure (Fig. 16). Figure 16A is the cross-section of the restored structure. From this, we excised single layers of myofilaments containing both myosin and actin filaments and only actin filaments, and created their projection maps in Figs. 16B and C, respectively. The graphs on the right side of the respective projections are the power spectra of the area boxed by green lines. The extent of reproduction of periodic structures varies from filament to filament. In Fig. 16B, the myosin filament boxed by green lines shows the clearest features of myosin heads. The power spectrum on the right shows 3 prominent peaks corresponding to the 1st, 2nd and 3rd reflections at d-spacings of 14.5, 7.2, and 4.8 nm, respectively. In the second myosin filament from the right, however, actin-like features are observed.

**Figure 16.**
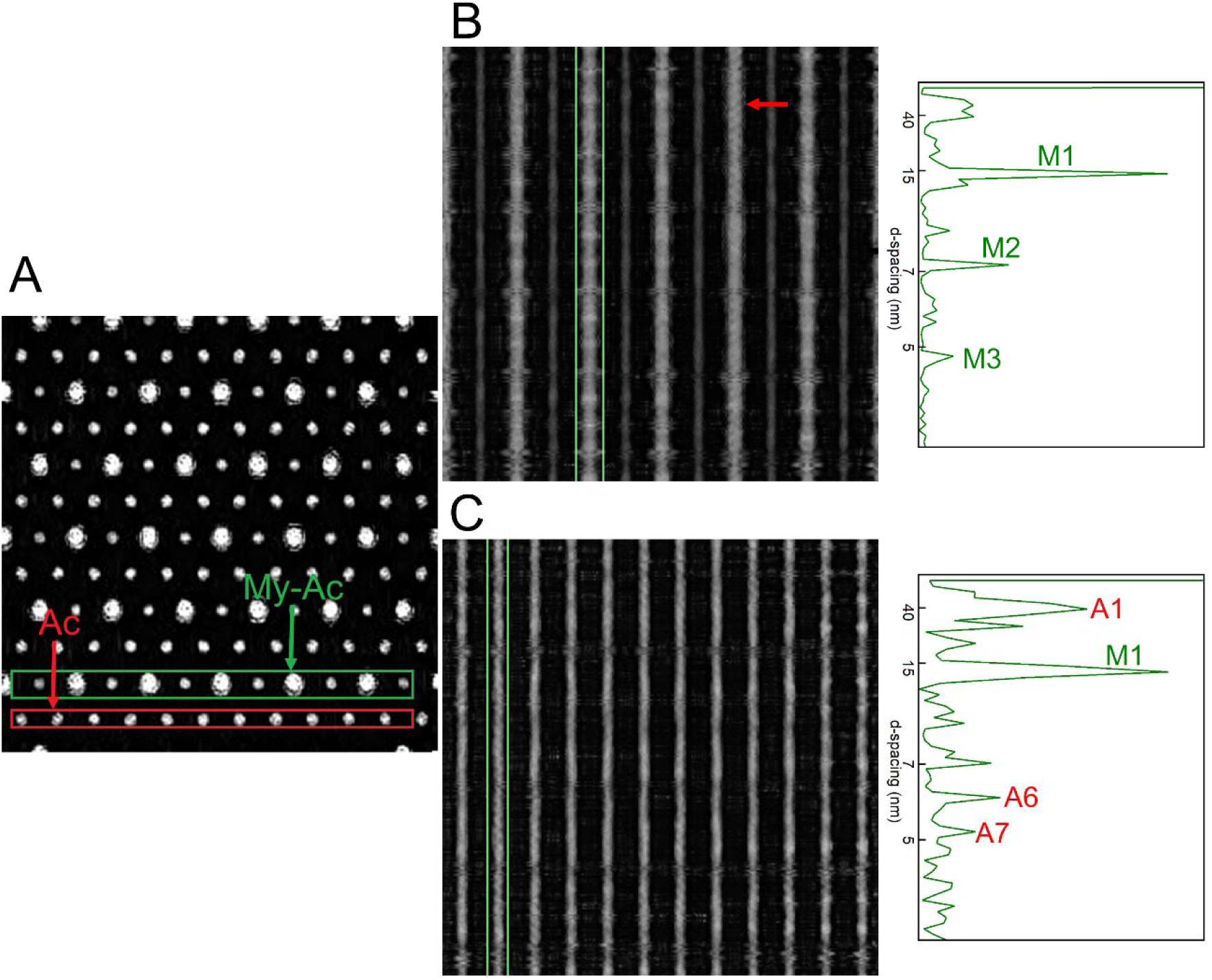
Density projection maps of the 3-D sarcomeric structure restored from the diffraction pattern of *Lethocerus* flight muscle fibers. (A) cross-section. (B) and (C) side views of single filament layers. (B) myosin-actin layer that contains both myosin and actin filaments (green box in A). The right graphs show the power spectra of the areas indicated by the green boxes in (B) and (C). M1, M2, and M3 correspond to the 1st, 2nd, and 3rd peaks originating from the 14.5-nm repeat of myosin heads. In (C), A1 corresponds to the 38.7-nm cross-over repeat of the actin double helix. A6 and A7 correspond to the left- and right-handed genetic helices of actin with pitches of 5.9 nm and 5.1 nm, respectively. Additionally, the M1 peak is observed. In (B), on the second myosin filament from the right, actin-like features are restored (red arrow). Likewise, in the rightmost two actin filaments in (C), myosin-like repeats are restored.

In Fig. 16C, the long-pitch periodicity of 38.7 nm is observed in most of the actin filaments, and the features of individual monomers are best reproduced in the filament boxed by green lines. In the power spectrum, 3 actin-based peaks are observed (marked A1, A6, and A7) at d-spacings of 38.7, 5.9, and 5.1 nm. However, a myosin peak (marked M1) is also observed. The myosin periodicity is visibly present in the two rightmost actin filaments in the projection. Therefore, the features of myosin and actin are generally, but not perfectly, separated in this reconstruction.

## Discussion

In this paper, we have demonstrated that the 3-D lattice structure of the sarcomere of insect flight muscle can be restored from a rotationally averaged 2-D structure factor using 3-D CDI, following optimization for 3-D FFT calculations. In the case of the in-silico experiment, the restoration was reasonably good, but even from an actually recorded diffraction pattern, a 3-D lattice structure was restored, although the quality of the diffraction pattern was much less favorable than in the in-silico experiment. The successful restoration is attributed partly to prior knowledge about the lattice structure and the helical symmetries of filaments in insect flight muscle, as documented by Tregear et al. (1998, 2004). We have previously shown that the 3-D structure of a fibrous material can be restored from its rotationally averaged 2-D structure factor through a completely different approach, i.e., a cylindrically averaged Patterson function (Iwamoto, 2021). This approach will work for a single-component system, but the sarcomeric structure of muscle contains two types of filaments, and its cylindrically averaged Patterson function will be too complex to solve. Therefore, 3-D CDI is better suited for the sarcomeric structure.

The tested size of the rotationally averaged structure factor is 256 x 256 pixels, and it is not a satisfactory size to capture all the features recorded in the real diffraction pattern. Experimentally, 1024 × 1024 pixels or larger is desirable to record all the fine structures of diffraction patterns from muscle, and a much larger size is needed to capture the ‘interference fringe’ (Reconditi et al., 2004; Huxley, 2004; Huxley et al., 2006a, b), which arises from the longer periodicity of myosin filament.

Increasing the size of the diffraction pattern and the 3-D workspace would improve the accuracy of structure restoration, but it would also increase processing time and memory consumption. With a size of 512 x 512 pixels, it would take 16 days to finish 500 iterations using our system. In our system, an array of 512 x 512 x 512 elements can be declared without an error, but a greater array may cause an out-of-memory exception. To address these issues, further optimization of both software and hardware is necessary. The use of graphics processing units (GPUs) may accelerate the calculations.

Besides the increase in the size of the 3-D workspace, the next challenge includes the application to time-resolved data. We have recorded time-resolved diffraction patterns from bumblebees during wing-beats at 5000 frames/s (Iwamoto & Yagi, 2013), and from demembranated muscle fiber specimens during quick stretch at 2000 frames/s (Iwamoto, 2016). In the diffraction patterns from live bees, the patterns from muscle fibers with different orientations are overlapping (dorsal longitudinal muscle and dorsoventral muscle). In the case of demembranated muscle fibers, the fibers can be aligned in parallel, making them better suited for reconstruction. If a time-resolved 3-D reconstruction is achieved, it is expected to reveal the motions of contractile proteins with a sub-millisecond time resolution. This may be achieved by repeating the present 3-D CDI calculations for individual frames. In this case, the present software programs can be used as they are. However, if a 4-D CDI calculation is conducted by regarding time as another spatial axis, the efficiency may be improved, analogous to the expansion from 2-D to 3-D CDI, which enabled the restoration of fibrous structures. Again, further optimization of both software and hardware is required to achieve this.

## Acknowledgements

This work was supported by JSPS Kakenhi Grant No. 19K06777 to H.I. The diffraction pattern shown in Fig. 1 was recorded under approval of SPring-8 Proposal Review Committee (proposal No. 2007A1190).

